# Phytop: A tool for visualizing and recognizing signals of incomplete lineage sorting and hybridization using species trees output from ASTRAL

**DOI:** 10.1101/2024.09.02.610893

**Authors:** Hongyun Shang, Kaihua Jia, Minjie Zhou, Hao Yang, Yongpeng Ma, Rengang Zhang

## Abstract

Incomplete lineage sorting (ILS) and introgression/hybridization (IH) is prevalent in nature and thus frequently result in discrepancies within phylogenetic tree topologies, leading to misinterpretation of phylogenomic data. Despite the availability of numerous tools for detecting ILS and IH among species, many of these tools are lacking effective visualization, or are time-consuming, or require prior predetermination. Here, we addressed these shortcomings by developing a fast-running, user-friendly tool called Phytop. By defining ILS and IH indices to quantify ILS and IH, this tool can detect the extent of ILS and IH among lineages with high reliability, and can visualize them based on the gene tree topology patterns constructed using ASTRAL. We tested Phytop extensively using both simulated and real data, and found that it enables users to quickly and conveniently estimate the extent of ILS and IH, thus clarifying the phylogenetic uncertainty. Phytop is available at https://github.com/zhangrengang/phytop and is expected to conveniently contribute to the intuitive inference of genetic relationships among lineages in future research.

## Introduction

Understanding the process of speciation is crucial for elucidating phylogenetic relationships, yet it is often complicated by phenomena such as incomplete lineage sorting (ILS) and introgression/hybridization (IH) [1–8]. These processes must be carefully considered, as they frequently result in conflicting phylogenetic tree topologies, posing significant challenges to researchers [9]. For example, the phylogenetic relationships among species within the Superrosids, a large clade of angiosperms, often exhibit inconsistencies across different datasets and methods, largely attributed to ILS [10, 11]. Similarly, introgression or hybridization, particularly in cases of certain species of hybrid origin, can lead to numerous conflicts between gene trees constructed for species and their parental lineages, as is observed in cases such as *Juglans regia* and *J. sigillata* [12], the Ericales [13], *Papaver* [14], *Fragaria × ananassa* [15], *Aegilops tauschii* [16], and *Triticum turgidum* [16].

With the growth of phylogenomics data, many tools have been developed to infer ILS or IH, including HyDe [17], MSCquartets [18], theta/Reticulation Index [6], Dsuite [19], BEAST [20], PhyloNet [21], PhyloNetworks [22] and BPP [23]. Two main classes of methods have been adopted by existing tools to detect hybridization. The D-statistic (also known as Patterson’s D or ABBA-BABA test) [24] is one of the most well-known hybridization detection methods. Based on the D-statistic, HyDe [17] and Dsuite [19] can be used to infer hybridization from summary statistics calculated from site patterns of population or species quartets. However, the accuracy of this method can be impacted at high levels of ILS or by ancient hybridization scenarios [25]. Another major way of detecting hybridization is to estimate a phylogenetic network for the detection of IH. Most of the software that adopts this approach, including NANUQ [26] in MSCquartets [18], InferNetwork_MPL [27] and InferNetwork_ML [28] in the PhyloNet [21] and SNaQ [29] in PhyloNetworks [22], uses likelihood methods or Bayesian inference, based on gene tree topologies or subtree topologies. This method is generally better suited to genomic datasets that contain multiple taxa but is computationally costly, and the information derived solely from gene tree topologies may not be sufficient to accurately distinguish the direction of gene flow [30, 31]. The full likelihood method makes comprehensive use of gene tree topologies and branch lengths, allowing for a more accurate identification of various gene flow scenarios, but it comes with an significantly increased computational burden [30, 32]. Methods using this approach include MCMC-SEQ [33] in PhyloNet [21], SpeciesNetwork in BEAST [20] and BBP [23]. Although the full likelihood method is a more efficient approach for the detection of gene flow, and performs better in distinguishing different introgression scenarios, its substantial computational load limits the application when dealing with a large number of taxa [30]. Thus, it is generally recommended to use a strategy that initially uses the fast but less accurate methods to come up with alternative hypotheses, and then to use the more accurate but slow methods (e.g. BPP) to test the hypotheses [30].

An additional problem is that none of the abovementioned tools provide effective visualization to intuitively display both ILS and IH among lineages, which discourages researchers from analyzing evolutionary networks in a more simplified manner. Tools like PhyParts [34] and DiscoVista [35] offer visualization of the gene tree discordance but do not provide quantifiable explanations. Furthermore, most of these tools require substantial programming skills and cannot quickly and conveniently provide quantitative estimates of ILS and IH, which is essential for subsequent analyses. Therefore, in practice, there is an urgent need for a fast-running tool that can quantify both ILS and IH and facilitate visualization to help optimize research steps. Such a tool would enable researchers to prioritize the exclusion of lineages with few phylogenetic conflicts, and then further analyze hard-to-resolve lineages using a combination of more robust methods, saving significant amounts of time and improving efficiency.

In this study, we developed Phytop, a simple and intuitive visualization tool that reflects the heterogeneity within phylogenetic species trees inferred using ASTRAL [36]. Phytop is freely available on github (https://github.com/zhangrengang/phytop). By visualizing the proportions and metrics of inferred topological structures, Phytop not only enables visualization of gene tree discordance along with each node of a species tree, but also provides interpretable measures of ILS and IH with high reliability. Even for phylogenetic trees comprising many lineages, Phytop can complete the computational process in a very short time, addressing the limitations of current tools. Moreover, Phytop, which visualizes the results of the popular tool ASTRAL for species tree inference [36], is easy to initiate and to understand, allowing for convenient, quick, and effective detection of the ILS and IH signals among lineages.

## Results

### Definition of the ILS and IH indices and expectations arising from these indices

We developed two indices for the visually assessment of ILS and IH signals in a species tree. We take a rooted species tree of three taxa as an example. The three taxa are the first child (or left child, L), the second child (or right child, R), and the sister group (S) (**Fig. 1a**). The rooted gene trees of the three taxa have at most three topologies: ((L, R), S), ((L, S), R) and ((S, R), L), with proportions of q1, q2 and q3 (q1 + q2 + q3 = 1), respectively (**Fig. 1b**). Under an ILS-only scenario (**Fig. 1a**), q2 = q3 is expected. When IH occurs with no ILS, for example an IH event from S to L (**Fig. 1a**), we expect q2 >> q3. For ease of understanding, we defined two indices to separately quantify the strength of ILS and IH signals. The first, named the “ILS index”, is scaled to a range of 0 to 100 % and reflects the strength of ILS. When the ILS index is at its maximum of 100 % and there is no IH, we expect that q1 = q2 = q3 = 100 %/3 (**Fig. 1c**). The second measure, abbreviated as the “IH index”, is essentially equivalent in concept to the previously defined inheritance probability (γ), gene flow rate (λ) or hybridization index [17, 37]. This index ranges from 0 to 50%. When the IH index reaches 50% and there is no ILS, q1 = q2 = 50% is expected (**Fig. 1d**). When both ILS and IH are present, the proportions of q1, q2, and q3 correspond to the different ILS index and IH index values, as shown in **Fig. 1e**. Under these expectations, the two indices can be therefore estimated through the patterns of q1, q2, and q3 (output from ASTRAL) and visualized using our Phytop tool. For the computational methods, please refer to the Methods.

**Figure 1.**
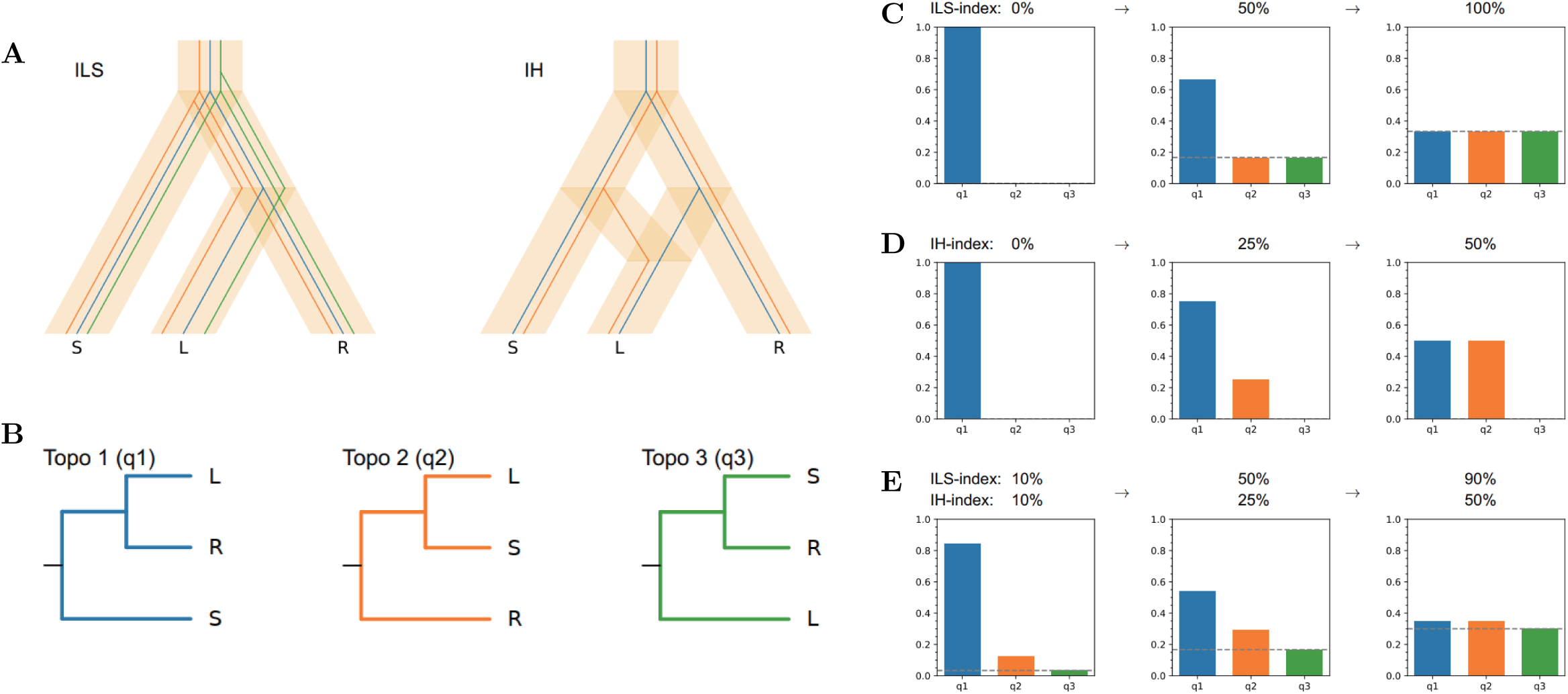
Theoretical expectations of the ILS and IH indices, taking the three species S, L, and R as an example. (a) The rooted species tree ([S, [L, R]]) and gene trees for ILS-only and IH-only scenarios. There are three and two gene tree topologies under ILS-only and IH-only scenarios, respectively. **(b)** The three possible topological structures of the gene trees, with the first topology identical to that of species tree. The proportions of the three topologies are denoted q1, q2 and q3. **(c)** Patterns of q1, q2 and q3 under an ILS-only scenario with different ILS index values. **(d)** Patterns of q1, q2 and q3 under an IH-only scenario with different IH index values. **(e)** Patterns of q1, q2 and q3 when both ILS and IH are present with different ILS index and IH index values.

### The performance of the indices in a simple hybridization model

We constructed simulated data to evaluate the reliability of the ILS and IH indices. In the simple hybridization model, lineage 1 served as the outgroup, and lineage 3 was a hybrid originating from lineages 2 and 4, as illustrated in **Fig. 2a** (For all the phylogenetic trees constructed by ASTRAL in this study, Phytop can be completed in just a few minutes). We set the ILS index to range from 0 to 1, with increments of 0.1, and set the IH index to range from 0 to 0.5, with increments of 0.05. For example, when we set the IH index to 0.4 and the ILS index to 0.368, we observed an IH index of 40.5 % and an ILS index of 36.1 % using Phytop (**Fig. 2b**), which was highly consistent with the expected values. For the full simulated data, the observed ILS index was essentially equal to the expected ILS index, with little fluctuation, and was essentially unaffected by variations in the IH index. The observed IH index was found to align for the most part with the expected IH index, although the fluctuation was large at high ILS levels and IH was sometimes even undetectable (**Fig. 2c**). For instance, when the ILS index was set to 1, the observed IH indices were zero across the board, and IH was thus completely undetectable (**Fig. 2c**). However, this is somewhat expected, as when ILS is particularly high, speciation is not complete, and there is no real inter-species IH. As the ILS levels increased, the variations in the observed IH index increased, and the detectability of the IH index decreased (**Fig. 2c-d**). Thus, with ILS levels increasing, a low IH index appears to be undetectable, and the undetectable boundary increases exponentially with increasing ILS levels (**Fig. 2d**). In addition, the error rates of the observed species tree topology increase, when either the ILS index or the IH index is high and close to their maximum values (**Fig. 2d-e**). However, this is also expected, because the three equivalent topologies are expected when ILS index reaches 100 %, and two equivalent topologies are expected when the IH index reaches 50 % (**Fig. 1c-d**), which lead to different topologies from the expected species tree topology. In summary, the observed IH index was found to be greatly affected by the ILS level, while the observed ILS index was almost totally unaffected by IH. The performances of the indices met our expectations using simulated data and suggest they are reliable.

**Figure 2.**
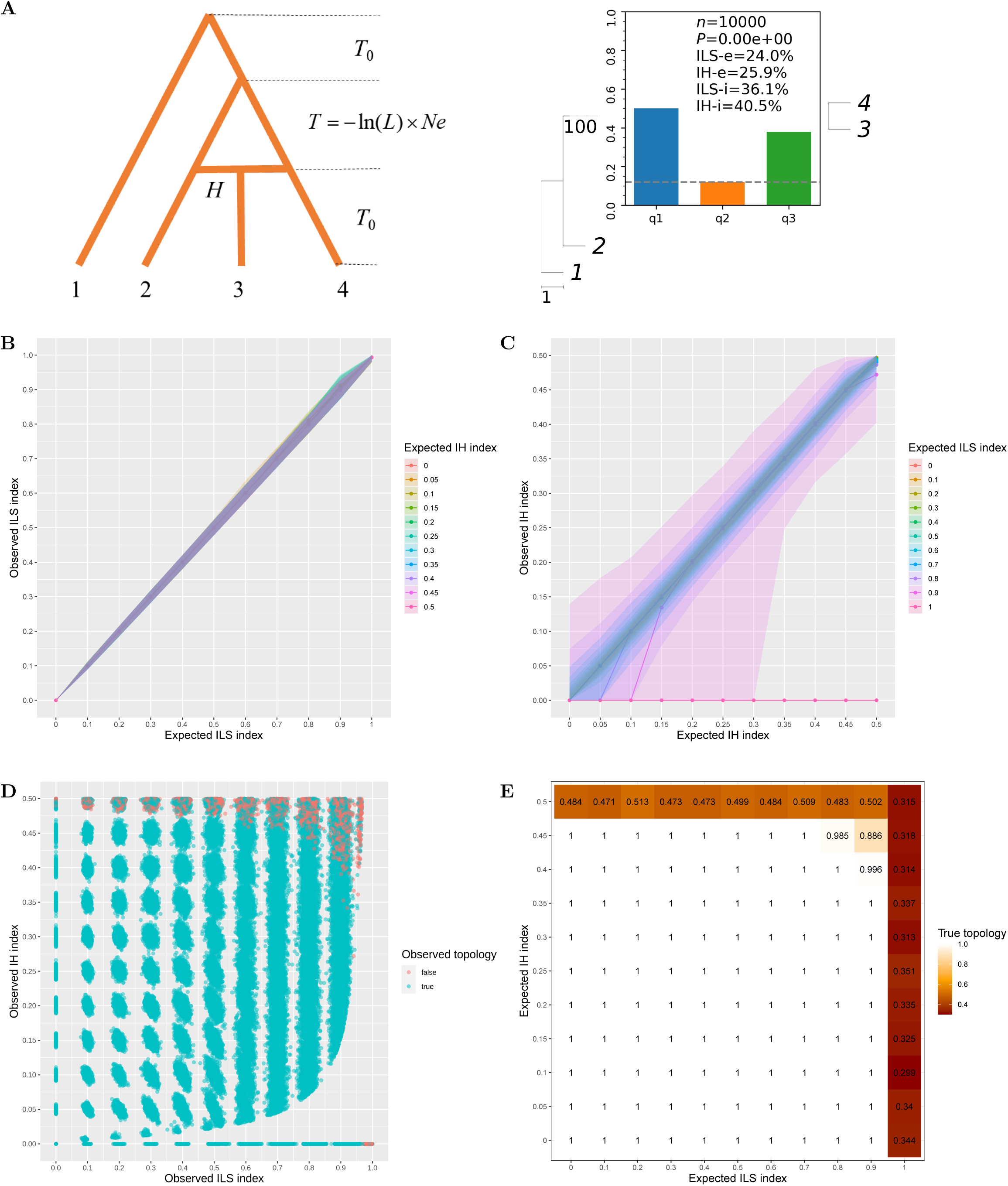
Evaluation of the IH and ILS indices in simulated data under a simple hybridization model. (a) We used fastsimcoal2 to simulate data for the simple hybridization model. In this model, lineage 3 is a hybrid resulting from hybridization between lineages 2 and 4 (left). *L* and *H* represents the ILS index and IH index set in the simulation, respectively. A more detailed description of the model parameters can be found in the Methods. An example of assessment of the degree of ILS and IH in the simulated data using the ILS and IH indices is shown (right). *n* represents the number of simulated gene trees, *P* is the p-value of the χ test to check whether the number of topologies q2 and q3 are equal, ILS-i and IH-i represent the calculated ILS and IH indices, respectively, and ILS-e and IH-e represent the proportion of gene tree topological incongruence that can be explained by ILS and IH, respectively. **(b)** The relationship between the expected ILS index and the observed ILS index under different settings of the IH index in simulated data. The shallows indicate 95 % CI of the observed ILS index. **(c)** The relationship between the expected IH index and the observed IH index under different settings of the ILS index in simulated data. The shallows indicate 95% CI of observed IH index. **(d)** The distribution of the observed ILS index and the observed IH index in all simulated data. Red dots represent unexpected topologies, while green dots represent expected topologies. **(e)** The proportions of observed true topologies under different settings of the ILS index and IH indices in simulated data.

### The performance of the IH and ILS indices in complex reticulation models

Although simple evolutionary models are seen in nature, complex reticulate models of evolution are perhaps more common in biological processes. We simulated several reticulate evolution models to investigate their effect on the gene tree frequency patterns and our two test indices, as shown in **Fig. 3**. For reticulation models 1-2 (with a single hybridization event), significant IH index values were observed in multiple nodes, although the hybridization branch was set for only one node (**Fig. 3a-b**). For reticulation models 3-4, where two hybridization events occur dependently, significant IH index values were also observed in multiple nodes (**Fig. 3c-d**), reflecting complexity and difference from the expected IH index value. None of the complex models resulted in an accurate estimation of ILS/IH indices. However, the detection of significant IH index value can reflect the presence of IH that needs to be further investigated using network-based methods, such as PhyloNet.

**Figure 3.**
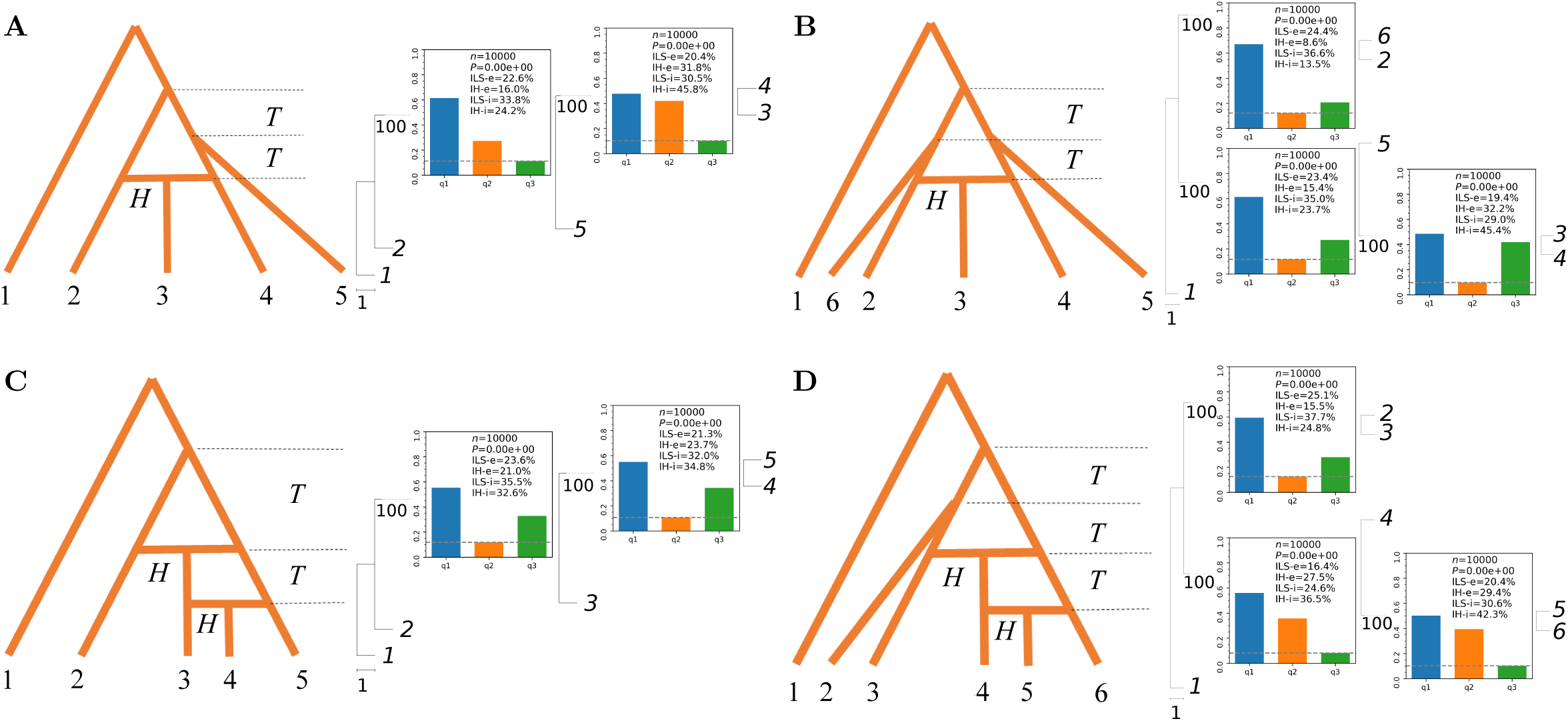
Evaluation of the IH and ILS indices in simulated data using complex reticulation models, with a single hybridization event in the reticulation models 1-2 (**a-b)** and with two successive hybridization events in the reticulation models 3-4 (**c-d**). The figures on the left show a schematic diagram of the model, and those on the right figure shows the phylogenetic relationships and indicators of the simulated data. All *H* and *T* branches are set to IH index = 0.4 and ILS index = 0.368, respectively.

### The performance of the IH and ILS indices in empirical data

We next evaluated Phytop using several previously reported cases of hybridization or introgression in *Juglans regia* and *J. sigillata* [12], the Ericales [13], *Aegilops tauschii* [16], and *Triticum turgidum* [16]. The ancestors of *J. regia* and *J. sigillata* are thought to have originated through hybridization between the American and Asian lineages [12]. We reconstructed the phylogenetic relationships among *J. regia*, *J. sigillata*, *J. mandshurica* (Asian lineage), and *J. nigra* (American lineage). We found that the ILS index at the node (*J. nigra*, (*J. regia*, *J. sigillata*)) was 48.1%, and the IH index, 44.2%, reflecting a potential hybridization event involving both the American and Asian lineages into the ancestors of *J. regia* + *J. sigillata* (**Fig. 4a**). This is consistent with the results from previous studies and further demonstrates the reliability of the ILS and IH indices in simple hybridization models.

**Figure 4.**
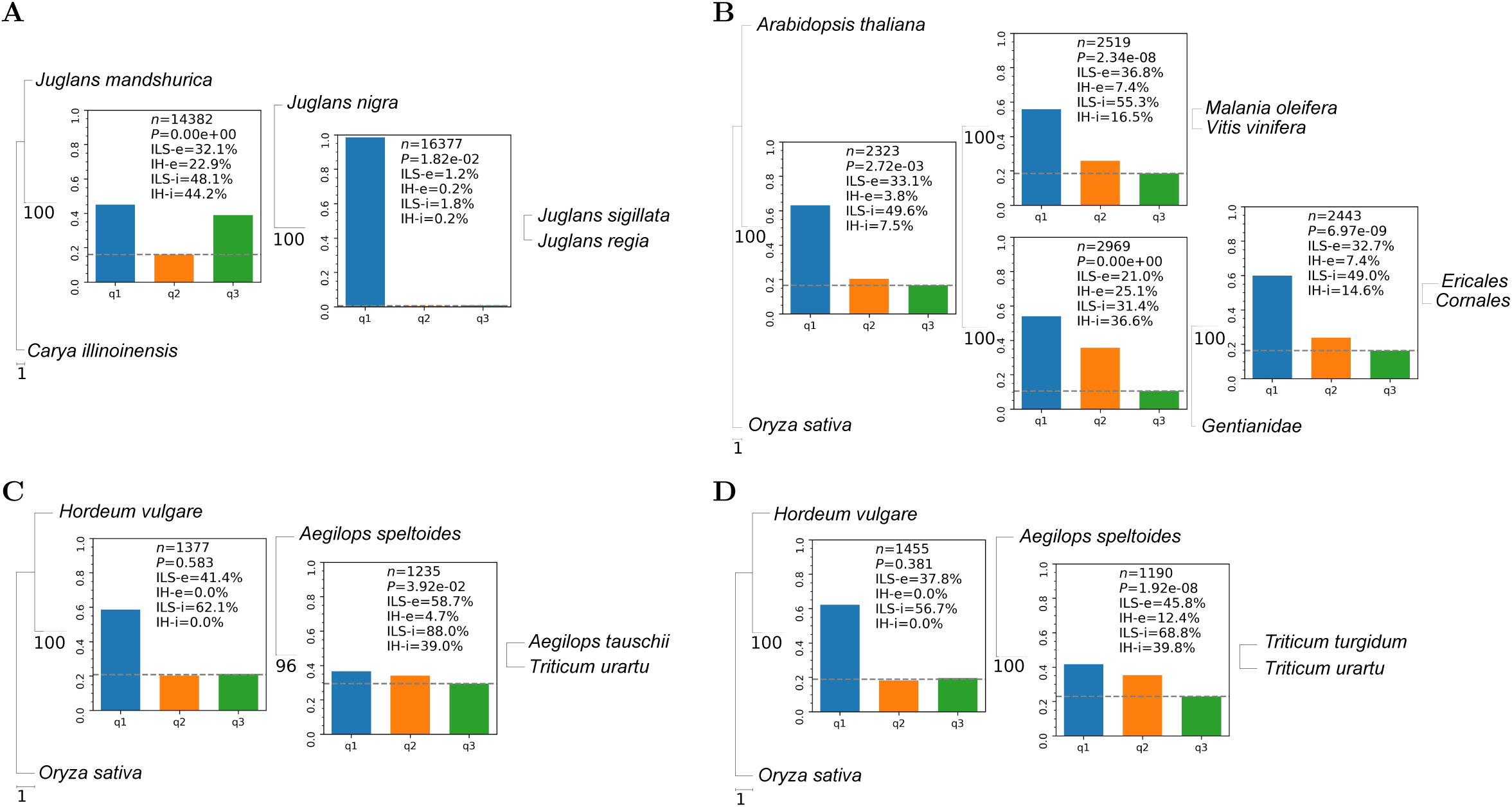
Evaluation of the ILS and IH indices in empirical cases of reticulate evolution. (a) Evaluation of the ILS and IH indices in the case of reticulate evolution within *Juglans*: *J. regia*, *J. sigillata*, *J. mandshurica* (Asian lineage), and *J. nigra* (American lineage). (b) Evaluation of the ILS and IH indices in the case of reticulate evolution among the Ericales, the Cornales and the Gentianidae. (c) Evaluation of the ILS and IH indices in the wheat D lineage (*Ae. tauschii*), and the A (*T. urartu*) and B (*Ae. speltoides*) lineages. (d) Evaluation of the ILS and IH indices in the allotetraploid emmer wheat (*T. turgidum*; AABB), and its progenitors: the A (*T. urartu*) and B (*Ae. speltoides*) lineages.

The Ericales are inferred to have originated from hybridization between the Cornales and the Gentianidae [13]. We selected genomic data from the Cornales, the Gentianidae, and the Ericales to reconstruct a phylogenetic species tree of these groups. The resulting IH index at the node ((Ericales, Cornales), Gentianidae) reached 18.5 %, which is essentially consistent with the degree of introgression (22.3 %) found in previous studies (**Fig. 4b**) [13]. This indicates potential gene introgression from the Gentianidae to the Ericales, supporting the previous hypothesis regarding a hybrid origin of the Ericales [13]. However, considering that there was also an IH signal at the adjacent node (the common ancestor of the Cornales, the Gentianidae, and the Ericales), a more complex reticulate model should be considered, according to our simulation of complex reticulate models (**Fig. 3**).

The wheat D lineage is hypothesized to have a hybrid origin resulting from hybridization between the A and B lineages [16]. Using *Oryza sativa* and *Hordeum vulgare* as outgroups, we reconstructed the phylogenetic relationships among *Aegilops tauschii* (D), *Ae. speltoides* (B), and *Triticum urartu* (A). We found that the IH index at the node (*Ae. speltoides,* (*Ae. tauschii*, *T. urartu*)) was 39.0%, which is consistent with the degree of gene introgression (39-57 %) estimated in previous studies (**Fig. 4c**) [16]. Furthermore, we found that the ILS index was 88%, indicating a strong level of ILS among the three lineages.

The allotetraploid emmer wheat (*Triticum turgidum*; AABB) is thought to have originated from the hybridization of the A and B lineages [16]. The phylogenetic relationships among *Ae. speltoides* (B), *T. urartu* (A), and *T. turgidum* were reconstructed (**Fig. 4d**). Similar to the case with the D lineage, we found that both the IH index and the ILS index at the node of (*Ae. speltoides,* (*T. urartu*, *T. turgidum*)) were at high levels (IH-i = 39.8%), which is consistent with the allotetraploid nature of *T. turgidum* potentially originating from the hybridization between *Ae. speltoides* and *T. urartu*.

These empirical cases suggest the reliability and accuracy of the IH index in detecting simple IH in real biological data. At the very least, the IH index is competitive with previously developed methods of inference.

### The performance of the ILS and IH indices in large biological datasets

We next explored whether ILS and IH signals were widely present in the major clade of plants (angiosperms and gymnosperms). Angiosperms, which form the most advanced class in the plant kingdom, are the most widely distributed group of modern plants [38–40]. We reconstructed species trees for several representative groups of angiosperms, including the early angiosperms, the core eudicots, the asterids, and the rosids.

In the early angiosperm groups, we found that ILS and IH signals were both widespread (**Fig. 5a**). We observed strong ILS and IH signals at the nodes that include branches such as *Aristolochia fimbriata*, the other Magnolianae, the Chloranthales, and the Ceratophyllales, where there was clear conflict in their phylogenetic relationships [41]. This suggests that the early diversification of the angiosperms may have involved complex reticulation. Similar to previous cases, the ILS and IH index are sensitive indicators that can reflect these factors effectively.

**Figure 5.**
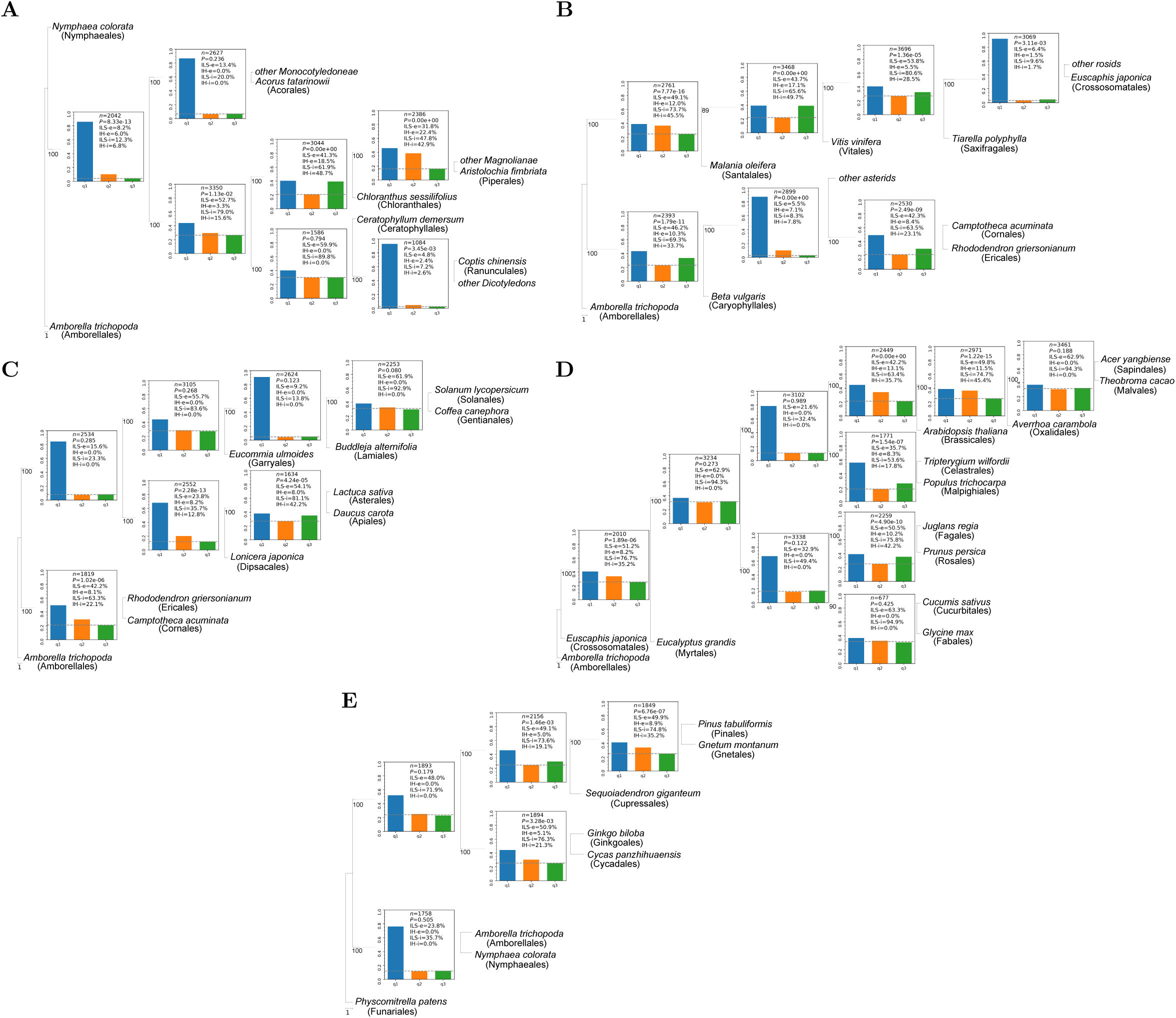
Detection of ILS and IH signals in representative groups of angiosperms and gymnosperms. Detection of the ILS and IH indices in the nodes in (a) early angiosperm groups, (b) early core eudicot groups, (c) the superasterids, (d) the superrosids, and (e) gymnosperm groups.

In the early core eudicot group of angiosperms, we found very strong ILS and IH signals among species, reflecting the severe ILS and IH events within the early diversification of core eudicots (**Fig. 5b**). In the superasterids group, ILS was also prevalent. In particularly, we found that the ILS index between the Solanales (*Solanum lycopersicum*), the Gentianales (*Coffea canephora*), and the Lamiales (*Buddleja alternifolia*) was as high as 92.9 %, although we detected no IH signals among the four species within the lamiids group (**Fig. 5c**). However, we found IH signals between species within the asterids, with the Dipsacales (*Lonicera japonica*) and the Asterales (*Lactuca sativa*) showing potential genetic introgression (IH-i = 42.2 %) towards the Apiales (*Daucus carota*). In the superrosids groups, we also found widespread evidence of ILS and IH (**Fig. 5d**). Our phylogenetic analysis of the superrosids groups showed significant divergence from previous studies where trees with high support were obtained using different datasets and methods (concatenation or coalescence), although previous studies have not been entirely consistent [10, 11]. In our study, it can be seen from the index that despite the high support, considerable ILS and IH signals were present in the species tree (**Fig. 5d**). For example, we identified strong ILS signals in the fabids, and strong IH (IH-i = 42.2%) signals among the Fagales, the Rosales, and the clade comprising the Fabales + Cucurbitales. Thus, we believe that widespread ILS and IH within the malvids and fabids clades are likely to be the reasons for the conflicts in species tree topologies observed in different studies [10, 11]. If the indices developed in this study are used for visualization, these issues can be observed and can be further addressed, helping to interpret the data and infer the correct phylogenetic relationships.

In gymnosperms, we also found widespread occurrence of both ILS and IH. All representative nodes of the gymnosperm phylogenetic tree showed IH signals, suggesting that complex reticulation events may have occurred among the main lineages of the gymnosperms (**Fig. 5e**), necessitating further research.

## Discussion

Understanding the history of life requires the establishment of well-resolved phylogenetic species trees, where the tree’s inner nodes with high support are considered to be the final topology [42]. In general, assessment of support for phylogenetic trees relies on statistical support for bipartitions, i.e. the nonparametric bootstrap proportion (BP) or the posterior probability (PP). Higher BP and PP values close to 100% reflect a low level of uncertainty in that node of the phylogenetic tree and are considered to be high support of that topology. However, with the development of sequencing technology in recent years, there has been an exponential increase in the amount of phylogenomic data produced, together with a decrease in random errors. This has led to high support often being obtained, but this does not necessarily truly reflect the heterogeneity within the data [43]. Furthermore, even where nodes are well supported, conflicting species tree topologies can be obtained for the same group in previous studies [9–11, 41, 43] and in this study (**Fig. 5**). ILS and IH are thought to be largely the cause of these phenomena [4, 44]. Here, we defined the ILS and IH indices to quantify the degree of ILS and IH which can in turn be another assessment indicator for the support of the phylogenetic tree. Based on simulated data, our rough guidelines are that a node with ILS < 30 % and IH index < 5 % can be considered as well-resolved with high confidence. By defining the ILS and IH indices, ILS and IH signals can be conveniently identified within phylogenetic species trees constructed by ASTRAL [36], which is a popular tool family in the phylogenomic era.

Under a simple hybridization model, using either simulated or real data, we were able to infer reticulate evolutionary relationships based on the ILS and the IH indices. Particularly in cases of simple hybrid origin, we found that the IH index obtained was highly consistent with results from previous empirical studies [12, 13, 16], which indicates the reliability of these indices in simple hybridization models. Although in complex reticulate evolutionary models, constrained by the framework of a binary tree, it can be difficult to accurately discern reticulate evolutionary relationships using the IH index, the IH signal can nevertheless inform us that reticulate evolution is present, and that further in-depth study within the network is necessary. This is a very important point: tree building, as a simple and effective method, is favored by researchers, but assessment of the potential presence of ILS/IH has often been neglected for a tree-based interpretation. Moreover, due to widespread ILS/IH, different species tree topologies are often obtained in different studies [10, 11], leading to conflict and confused interpretations of phylogenetic relationships. The visualization made possible by Phytop could provide a straightforward evaluation for the uncertain or hard-to-resolve nodes (such as the (Apiales, Asterales) node with a high ILS or IH in Figure 5c) of species trees.

We analyzed the ILS and IH signals in the major groups of angiosperms and gymnosperms, and found that ILS and/or IH signals were widely present across a large number of groups. With the functionality provided by Phytop, researchers can quickly predict ILS/IH between species from phylogenetic trees constructed in ASTRAL [36] and can adopt a divide-and-conquer strategy for the inference of phylogenetic networks, Well-resolved nodes with both high support and low ILS and IH indices can be first excluded, and then researchers can focus on analyzing the phylogenetic networks of species involved in inner nodes with high ILS and/or IH indices. This is undoubtedly very important because this strategy greatly reduces the complexity and time-consuming processes in the operation of existing phylogenetic network analysis tools (such as PhyloNetworks [22] and BPP [23]), and is necessary for further elucidating the evolutionary relationships among a large number of taxa. The Phytop method provides simple and effective visual identification for ILS/IH, allowing researchers to interpret the uncertainty in species tree topologies fully and simply, and guiding researchers to further transition to a network framework. Moreover, Phytop is easy to operate and requires only a short run time, meaning that it is easily mastered by users.

## Methods

### Definition of ILS index and IH index in theory

Under the multispecies coalescent (MSC) model (only ILS), the two minor topologies (q2 and q3) have equal probability *e*^−^ *^t^* / 3, where *t* is the coalescent time [9]. We define the ILS index as *e*^−^*^t^*, so that when *t* = 0, it reaches 100% and the three topologies have equal probability (1/3), The ILS index drops to < 1% when *t* > 4.6. This index is more simple to understand than the coalescent time *t*. Given an ILS index value (denoted as *L*), the probability of a minor topology is *L*/3 and the coalescent time is *t* = − ln(*L*) . This index can therefore reflect the strength (0–100%) of ILS and the minor topology frequencies (0–0.33) of gene tree discordance.

When asymmetrical introgression/hybridization (IH) occurs, we define IH index as the proportion contributed by the minor donor lineage. Given an IH index value (denoted as *H*), a lineage is considered to be admixed by *H* of one parental lineage and 1−*H* of the other parental lineage. This index is also simple to understand, reflecting the proportion (0–50%) of the genome inherited from the minor parental lineage.

### Estimation of ILS index and IH indices

Given the frequencies *q*_1_, *q*_2_ and *q*_3_ (let 1 ≥ *q*_1_ ≥ *q*_2_ ≥ *q*_3_ ≥ 0; *q*_1_ + *q*_2_ + *q*_3_ = 1) of the three topologies in the species tree inferred by ASTRAL, we use a chi-square test to test the goodness-of-fit for the multi-species coalescent model [9]. If the *P* value is greater than 0.05, indicating that there is no significant difference between the observed and expected topologies, we say the topology frequencies can be well explained by only ILS (ILS can explain all the *q*_2_ + *q*_3_ discordant gene trees). In this case, we use (*q*_2_ + *q*_3_) / 2 to represent the minor topology frequencies, and the ILS index is estimated using the formula:

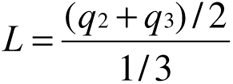

Otherwise (*P* ≤ 0.05), we say the smallest *q*_3_ can be explained by ILS (ILS can explain 2 × *q*_3_ discordant gene trees) and the imbalance between *q*_2_ and *q*_3_ can be explained by IH (IH can explain *q*_2_ −*q*_3_ discordant gene trees). Thus, we use *q*_3_ to represent the minor topology frequencies from ILS, and estimate ILS index using the formula:

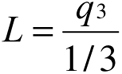

Furthermore, ILS itself is expected to produce three topologies with equal probability (*q*_3_). Thus, with *q*_3_ all excluded, the IH index is estimated using the formula:

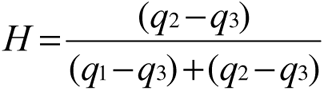

Note that other factors, such as horizontal gene transfer (HGT) and gene duplication and loss (GDL), are not addressed in our assumption. Nevertheless, the symmetries in gene tree distributions can be addressed using the ILS index, and the asymmetries can be addressed using the IH index. In addition, it is worth noting that the two indices are calculated under a simple hybridization scenario (Fig. 2a) and may not be accurate for other scenarios (e.g., Fig. 3).

### Simulation data

We simulated a mixed (ILS + IH) model using fastsimcoal2 [45]. Population effective sizes (*Ne*) and mutation rate were fixed to 1e5 and 1e-8, respectively; T0 = 1e6, the substitution rate =1e-8, the recombination rate = 0, the locus length of was set to 2000bp; the IH index was set from 0 to 50% with a step of 5 %, and the ILS index was set from 0 to 100% with a step of 10%, producing 121 (11×11) combinations. The generation time (*T*) of internal branches was converted from the ILS index (*L*) using formula: *T* = −ln(*L*) × *Ne* . Each scenario was simulated (‘fsc27 -n 10000 -T’ at each time) to generate 10,000 gene trees. Then, the species tree was inferred using ASTRAL-Pro2 [36] and the two indices (i.e. ILS index and IH index) were calculated based on the observations of gene tree frequencies. The above procedure was replicated 1,000 times independently to generate percentile-based 95% confidence intervals (CI) for the two indices.

### Real data

Four real biological datasets were used in this study:

1. The wheat data: Marcussen et al. [16] revealed a hybrid origin of the D lineage from the A and B lineages in the wheat genome. The allotetraploid AABB lineage is thought to have originated from the hybridization of the A and B lineages. Here we used the genomic data from *T. urartu* (AA), *Aegilops speltoides* (a close relative of BB), *T. turgidum* (AABB) and *Ae. tauschii* (DD) to test these inferences. *Hordeum vulgare* and *Oryza sativa* were used as the outgroups.
2. The walnut data: Zhang et al. [12] demonstrated that *Juglans regia* (and its landrace *J. sigillata*) arose as a hybrid between the American and the Asian lineages. Here we used the genomic data from *J. regia*, *J. sigillata*, *J. mandshurica* (Asian lineage) and *J. nigra* (American lineage) to test this inference. *Carya illinoinensis* was used as the outgroup.
3. The Ericales data: Stull et al. [13] suggested the possibility that the Ericales was a reticulate lineage resulting from hybridization between ancestors of the Cornales (78%) and the Gentianidae (22%). Here we used the genomic data from the Cornales, the Ericales, and the Gentianidae from our previous study [46] to test this inference. *Arabidopsis thaliana*, *Malania oleifera*, *Vitis vinifera*, and *Oryza sativa* were used as the outgroups.
4. The angiosperms data: In addition to the above well-documented data, we also used genomic data from the angiosperms to test our method. This dataset included 45 orders and 45 families of angiosperms, as well as two outgroups (*Amborella trichopoda*, and *Nymphaea colorata*).
5. The gymnosperms data: We selected representative gymnosperms to construct a species tree with *Physcomitrella patens* as the outgroup.

### Inferring species tree from biological data with ASTRAL

Orthogroups were inferred using Orthofinder [47] for the above real data. MCScanX [48] was used to extract gene collinearity. Single-copy gene families were extracted from the collinearity results, allowing up to 50 % missing taxa. Protein sequences in each family were aligned using MAFFT [49] and the alignments were converted into codon alignments using PAL2NAL [50]. The alignments were trimmed using trimAl, with the parameter ‘-automated1’ that is optimized for Maximum Likelihood (ML) phylogenetic tree reconstruction [51]. Then the ML phylogenetic trees were reconstructed by using the software IQ-TREE [52] with the automatically-selected best-fit model [53] and 1000 bootstrap replicates [54]. The gene trees were then re-rooted using the Newick utilities [55]. Based on the single-copy gene family trees, ASTRAL-Hybid [56] was used to infer the species tree with quartet supports (i.e. counts of the three topologies, or q1, q2 and q3) which is weighted by gene tree uncertainty.

### Visualizing the species tree

We visualized the frequencies of the three topologies using bar charts and integrated the species tree illustration with the Python programming library ete3 [57]. Two indices, the ILS index and the IH index, were calculated (see above) to quantify the strength of incomplete lineage sorting (ILS) and introgression/hybridization (IH) signals. These steps were done with Phytop (https://github.com/zhangrengang/phytop) and only take a few minutes for a tree with dozens of species.

## Acknowledgements

This study was financial support from Natural Science Foundation of China (32471734), CAS “Light of West China” Program, Yunnan Provincial Science and Technology Mission (202404BI090014). We thank for Jane Marczewski’s assistance with English language editing.

## Code availability

The code for Phytop is freely available on GitHub (https://github.com/zhangrengang/phytop).

## Conflict of interests

The authors declare no conflicts of interest.

## Author contributions

R. Z. and Y. M. conceived and designed the study. R. Z. performed software development. H. S., and R. Z. wrote the manuscript. R. Z. and H. S completed the data analysis and prepared the figures. R. Z., Y. M. and K. J. revised the manuscript. M. Z. and H. Y. tested the software and helped with data analysis.

## Notes

### Competing Interest Statement

The authors have declared no competing interest.

